# Learning reshapes the hippocampal representation hierarchy

**DOI:** 10.1101/2024.08.21.608911

**Authors:** Heloisa S. C. Chiossi, Michele Nardin, Gašper Tkačik, Jozsef L. Csicsvari

**Affiliations:** Institute of Science and Technology Austria, AT-3400 Klosterneuburg, Austria; Janelia Research Campus, Howard Hughes Medical Institute, Ashburn, VA 20147, USA

**Keywords:** hippocampus, CA1, representation, learning, hierarchy

## Abstract

A key feature of biological and artificial neural networks is the progressive refinement of their neural representations with experience. In neuroscience, this fact has inspired several recent studies in sensory and motor systems. However, less is known about how higher associational cortical areas, such as the hippocampus, modify representations throughout the learning of complex tasks. Here we focus on associative learning, a process that requires forming a connection between the representations of different variables for appropriate behavioral response. We trained rats in a spatial-context associative task and monitored hippocampal neural activity throughout the entire learning period, over several days. This allowed us to assess changes in the representations of context, movement direction and position, as well as their relationship to behavior. We identified a hierarchical representational structure in the encoding of these three task variables that was preserved throughout learning. Nevertheless, we also observed changes at the lower levels of the hierarchy where context was encoded. These changes were local in neural activity space and restricted to physical positions where context identification was necessary for correct decision making, supporting better context decoding and contextual code compression. Our results demonstrate that the hippocampal code not only accommodates hierarchical relationships between different variables but also enables efficient learning through minimal changes in neural activity space. Beyond the hippocampus, our work reveals a representation learning mechanism that might be implemented in other biological and artificial networks performing similar tasks.

## Introduction

Recent years have witnessed a growing interest in representation learning, both in biological and artificial neural networks. For example, representations in the early areas of the visual ventral stream are known to match simple features of the visual environment (1–4), and to reshape as a result of learning new input statistics (5–7). Recently, these representations have been shown to bear a close similarity to representations observed in convolutional neural networks trained for image classification (8–11). Similarly, motor learning can alter representations in the output layers of the mammalian motor system in a manner analogous to neural networks trained on similar tasks (12, 13). These developments have been central to the study of neural network dynamics (12, 14, 15) as well as to the development of neuroprosthetics (16, 17). Higher associational brain areas, such as the hippocampus, however, feature more abstract, mixed responses to environmental variables (18–22). An understanding of how these areas form their representations during learning could shed light on the network mechanisms that underpin behaviors beyond sensory classification or motor control.

One form of learning that still challenges artificial networks is associative learning (23). It involves the integration of newly acquired information with previously accumulated knowledge through the emergence or update of neural representations. In biological networks, neural recordings during behavior can help us identify strategies used by the brain to form representations during associative learning. Creating associations is extremely relevant for survival; e.g., an animal looking for food may act differently during day or night, guided by previous experiences with different predators and food sources associated with those two contexts. First, animals need to learn the representation of the local environment, to be able to navigate to behaviorally relevant locations such as rewards. Second, they need to establish an association between reward locations and the contextual cues that determine them. The hippocampus is a key area for both learning tasks: it represents variables such as place, together with other behaviorally relevant environmental cues (24–28), and is essential for establishing memory and context associations (for a review, see (29)).

In the hippocampus, spatial representations are formed within a few minutes, and these remain relatively stable over multiple days (30). Associative learning, however, can take much longer, typically on the scale of days or weeks for rodents (31–33). It is unclear how the representation of associative learning-related variables changes during this period. Nevertheless, it is known that single hippocampal neurons can be selective for multiple variables (18–22), a feature that increases the code’s repertoire of possible readout functions. This, however, implies that neural representations of single variables are distributed over multiple dimensions in neural activity space (34), with their structure reflecting the tasks an animal has been trained on (35). As a consequence, even key signatures of learning might only be detectable at the population scale.

Understanding how associative learning alters task-related representations in the hippocampal neural population could help elucidate general neural network mechanisms employed in complex task learning. To study this, we monitored, over several days, how representations in the hippocampal CA1 pyramidal cell population reorganized as naive animals learned a goal-oriented contextual task (Figure 1). We detected a hierarchy in the representation of different variables at the population level (20, 21, 36), which was maintained throughout the entire learning process. Changes in the population representations during learning were constrained to the bottom of the hierarchy, involving local adjustments that correlated with behavioral performance. We demonstrated that this reorganization was sufficient to enable better decoding of task-relevant features downstream, in addition to allowing for compression of task-relevant information into a few dimensions of high neural activity variance.

**Figure 1.**
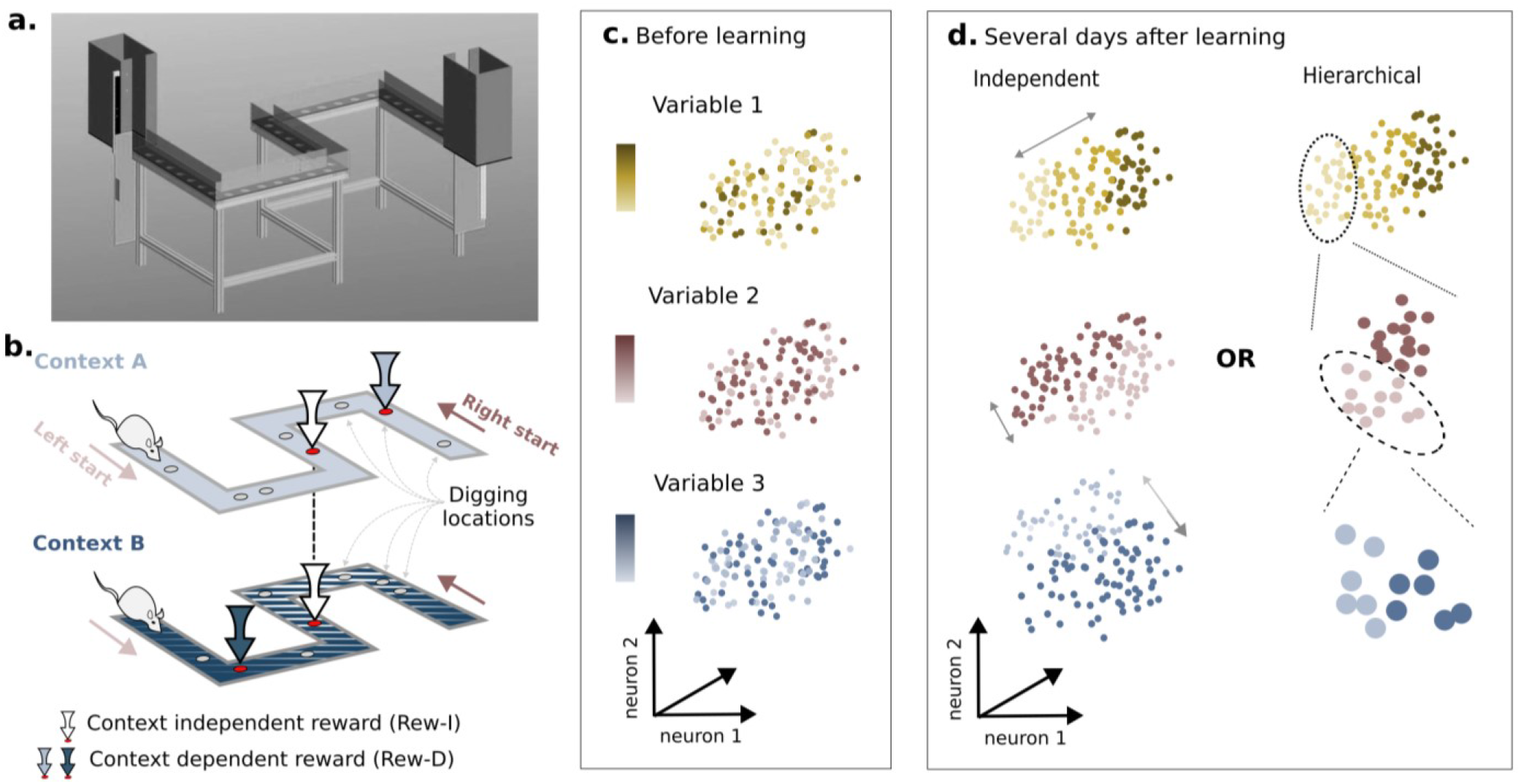
Experimental setup and hypothesis. **a)** Illustration of the experimental maze: A 360cm track (80cm arms overlapping by 10cm at the corners), containing two start/end boxes and 5 arms arranged into an S-shape, each containing 7 wells which can be filled with sand. **b)** Illustration of the task: Within the same physical maze, the color and texture of the flooring could be changed to indicate a new context. In each of the two contexts, 8 wells were made available to the animal, two of which contained a food reward. One well location was common to both contexts (“context independent reward”, or Rew-I), whilst the second reward depended on the context yet was not indicated by any direct sensory cue (“context dependent reward”, Rew-D). c-d) Colors indicate different task variables (e.g. position, direction, context) and different shadings indicate different values these variables can take. **c)** Before learning, task variables likely do not have a clear representation in neural activity space. **d)** We consider two possible coding schemes that can emerge during learning: in the “independent” scheme (left), variables are encoded in orthogonal directions of the high-dimensional neural activity space; in the “hierarchical” scheme (right), fixing the value of a variable at the top of the hierarchy (conditioning) reveals separable organization of neural activity for a variable encoded at a subordinate hierarchical level.

## Results

### Experimental paradigm for associative spatial learning

We designed a goal-oriented spatial task where context specified which position was rewarded. We built a 3.6-meter-long S-shaped linear track (Figures 1a and S1a) where visual and tactile cues could be changed between trials, thereby establishing the trial context (denoted A or B). The track contained multiple sand-filled wells in which a food reward could be hidden. On each trial, a piece of food reward was buried in two of the wells: one food-bearing well remained the same across both contexts (“context-independent reward” or Rew-I), while the position of the second food-bearing well differed between contexts (“context-dependent reward” or Rew-D) (Figure 1b). For a given animal, reward locations remained fixed between days, and wells at which an animal engaged in digging indicated its memory for the reward location.

On each training day, animals ran a fixed number of trials (40 trials). Trials started randomly from either end of the track, allowing us to sample both movement directions equally (left or right); trials finished when the animal entered the box opposite to their start location. On a given trial, animals were allowed to dig only two wells – the remaining wells were covered after the choices were made – and the trial was considered correct if they dug exactly the two rewarded locations. We opted for a digging task because digging is a highly motivated behavior: despite the food being placed at similar depths in all trials, animals had to engage in digging for a few seconds up to a few minutes before finding the food (Figure S1d), which made decisions costly for the animals and incentivized learning.

At a given session, an animal ran an equal number of trials from each combination of context and trial direction – which we refer to as trial category – and all positions were visited in all trials. We could thus track three task variables throughout the learning: position, context, and movement direction. The association between reward position and context was central to task performance, whilst associations between reward position and direction were not; these had to be ignored for correct allocentric navigation to the goals. Therefore, our experimental setup allowed us to monitor variables with different significance for task performance throughout the entire task learning period.

### Task-solving strategies change with learning

In 3-5 days, all animals (n=5) reached a task performance of >80% correct trials (Figure 2a). We identified two learning stages: in the first stage, animals identified the three wells that could be rewarded (1 Rew-I and 2 Rew-D) in as little as the first 7-8 trials. From then on, they dug whichever two of those three wells they encountered first, rather than using contextual information to make a choice. We call this an “eager” task-solving strategy. In the second stage – after a few days – the choice was correctly based on the context, so we call it a “contextual” strategy. We quantified the use of each strategy by measuring the number of trials in a given day in which an animal’s choice was consistent with one or the other (Figure 2b).

**Figure 2.**
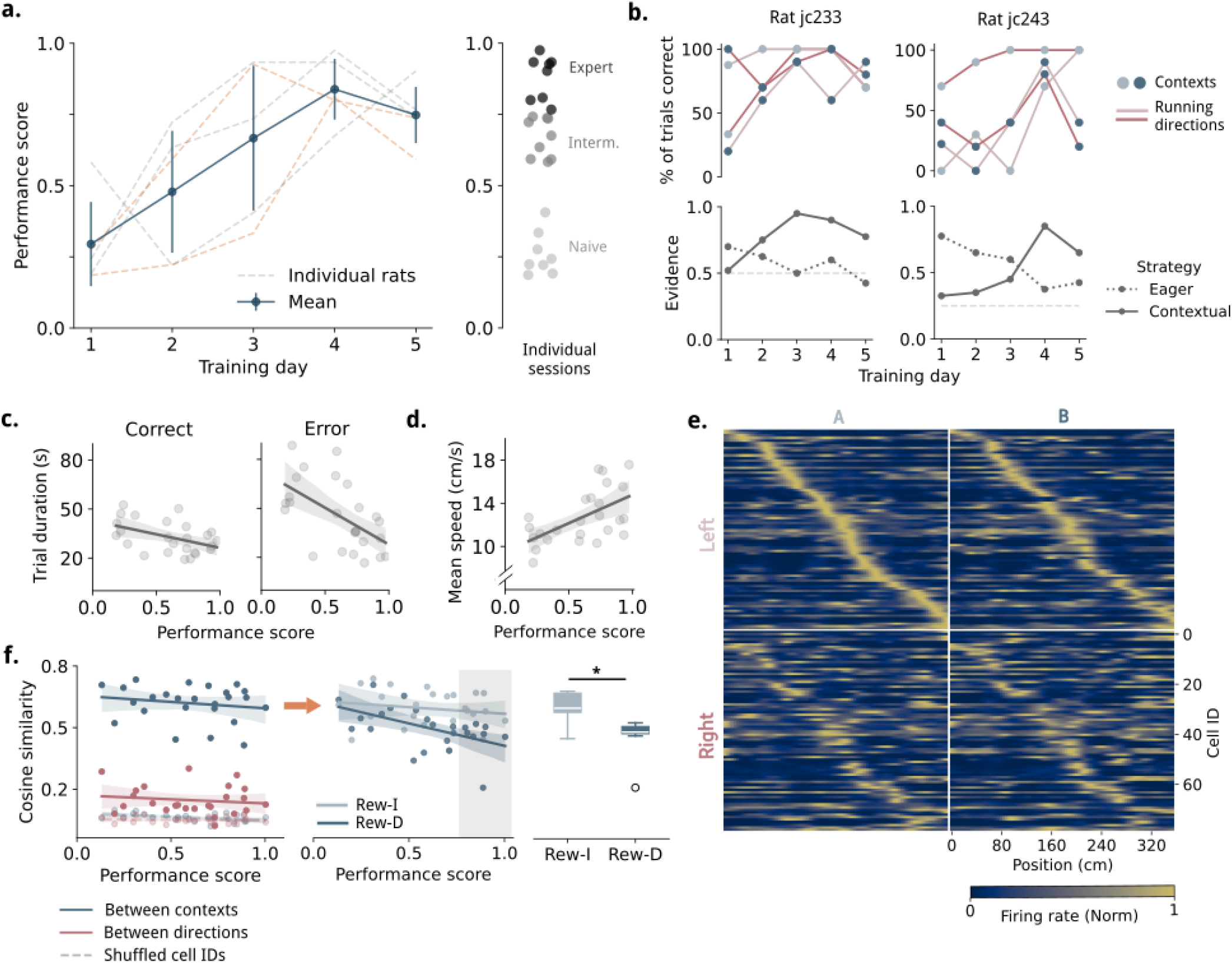
Behavior and neural population tuning. **a)** Left: Training increases animals’ behavior performance score over multiple days. Performance on a given trial was 1 if an animal dug exactly the two rewards appropriate for the given context and nothing else (“correct trial”). Blue line indicates mean +- SEM for all animals (n=5); dashed lines indicate the average daily performance of individual animals. In yellow are highlighted the examples shown in panel (b). Right: Sessions were assigned to three quantiles of performance: naive (n=8), intermediate (n=9) and expert (n=8). **b)** Top: Percentage of correct trials for each category (= combination of context and direction) separately, for two example animals. Bottom: Evidence that the animal was using a specific task-solving strategy. “Eager” refers to digging the first two of the three rewardable holes, given the trial direction. “Contextual” refers to digging exactly the two holes rewarded, given the trial context. The horizontal gray lines indicate the expected evidence for eager strategy given a perfect contextual strategy; this value is specific to the reward configuration each animal experienced in the maze. **c)** Left: Trial duration, excluding digging periods, versus the daily performance score for correct trials. p < 7e^-3^ R = -0.44. Right: Same, but for error trials. p < 0.03 R = -0.54 needs to be changed. **d)** Mean speed increases with performance score (n=25 sessions). p < 3e^-3^ R = 0.57. **e)** Normalized firing rate maps of all pyramidal neurons on an example recording day, in all 4 trial categories (rows: left/right direction; columns: A/B context). Neurons were ordered according to their peak firing position in left direction + context A trials (upper left category). **f)** Left: Average cosine similarity of population vectors calculated per position, between rate maps of opposite directions / same context (pink) or opposite contexts / same direction (blue), averaged over all positions. Calculated from the GLM speed-free rate map. Dashed lines indicate the same calculation using shuffled cell IDs (see Methods). All correlations with performance score were n.s.. Middle: Similarity values between contexts as a function of performance only around the specific reward location (within 12 cm radius from reward center). Rew-I n.s.; Rew-D R = -0.65 p < 0.01. Context is most dissimilar between the two reward locations in expert sessions (shaded gray area), as shown also in the right panel. Right: Cosine similarity between contexts in expert sessions only, around reward locations. Rew-I vs Rew-D in expert sessions p < 0.01 Wilcoxon test. Regression values calculated with partial linear regression controlling for the number of pyramidal neurons per session and animal IDs. In all regressions, shaded areas indicate a 95% confidence interval.

However, eager choices sometimes matched the correct contextual choice. When the Rew-I was located between Rew-Ds in the linear track, the eager strategy could yield up to 50% of trials correct by chance, but only 25% in configurations where Rew-Ds were located after each other. Since reward locations were randomly assigned, this meant that the eager strategy yielded more correct trials for some of the animals. To correct for this, we calculated the proportion of trials that were unambiguously eager trials and used it to estimate the proportion of eager trials from those where both strategies were possible (see Methods). This resulted in a score that was close to zero when the evidence was greater for the eager strategy and increasingly positive up to a value of one as the animal approached a perfect contextual strategy. While this performance score was based solely on animal choices, it was nevertheless highly correlated with a number of other behavioral measurements. Both correct and error trials became shorter in duration as the performance improved, pointing to more certainty in the animals’ behavior (Figure 2c). Also, animals with better performance moved faster (Figure 2d), spent less time at the incorrect reward location when they made mistakes (Figure S1e) and were more likely to refrain from digging when running past the incorrect reward (Figure S1c). We divided the sessions into three equally sized quantiles of performance, which we denote *naive, intermediate*, and *expert* (Figure 2a and S1b). Those were used in later analyses in which performance was treated as a discrete variable.

Our behavior characterization allowed us to relate changes in neural representations directly to behavioral performance rather than to training time (in terms of days). This led to more interpretable results by discounting individual differences in training trajectory over time and thus yielded a higher correlation between behavior and neural representation measures.

### Learning drives local divergence of context representations

Once the behavior was characterized, it was possible to investigate how the hippocampal population activity represented the task as the animals learned. We simultaneously recorded the activity of multiple neurons (n=1725 putative units in total, mean of 69 ± 25 units per day per animal) from the dorsal CA1 region of the rats using movable tetrodes, across all five days of task learning. On average, 80 ± 10% (mean ± STD) of cells recorded in a given day were classified as putative pyramidal cells, which were used for the analyses presented below.

A large proportion of hippocampal pyramidal neurons displayed strong spatial selectivity, and the recorded neural population on any day was sufficient to tile the entire environment (examples in Figure 2e and S2a). Moreover, these cells were sensitive to changes in movement direction and context, with a larger effect of the former over the latter (examples in Figure 2e and S2a). We also expected movement speed to be a modulator of the hippocampal activity (28, 37), but it was inherent to our behavior task that speed was unevenly sampled. To guarantee that our analyses were not biased by correlations of other variables with speed during different trials, we performed two precautions: (i) we only considered running periods (> 3cm/s) for the analyses; (ii) we excluded the speed contribution to the firing rate at each position and trial by fitting a Generalised Linear Model (GLM) (38, 39) for each cell and jointly estimating the contribution of speed and position to each cell’s mean firing rate. This allowed us to generate “speed-equalized” rate maps for each cell, which we used in subsequent analysis (Methods, Figure S3). Nonetheless, the analyses we report here yielded qualitatively similar results when calculated on the original or speed-equalized data.

Using the per-session speed-equalized rate maps, we extracted the firing rate of each cell in each position (discretized into 90 bins of 4cm), averaged over all trials of a category, to form a population vector (PV). Each PV was associated with labels for context (A or B), movement direction (left or right) and position. To measure selectivity to context or direction, for each maze position, we calculated the cosine similarity between PVs from each condition. The average similarity over all positions between different contexts was generally higher than between different movement directions. Surprisingly, there was no significant change in similarity with increasing behavioral performance (Figure 2f, left).

We, therefore, wondered if changes in representation were more local and consequently obfuscated by averaging across positions. We focused specifically on positions where context had behavioral significance, i.e., on Rew-I and Rew-D maze positions. At Rew-Ds, PVs of different contexts became increasingly dissimilar as performance improved, but not at Rew-I (Figure 2f, right). This indicated that the hippocampus represented context better only where it was relevant for decision-making. Moreover, the overall population activity differed between these positions in terms of population firing rate gain. A gain value of 1 indicates that the summed activity of all neurons at a certain location was equal to the average population activity across all locations. Based on previous reports, we expected that all reward locations would exhibit higher population activity than unrewarded locations (40, 41) and that familiarity would lead to a decrease in this effect over time (42). In contrast to this expectation, at Rew-I, the population activity gain remained at 1 despite it being a rewarded location; at Rew-D the population activity was above average, regardless of whether the position was rewarded in the current trial or not (Figure S2b). We also did not observe a reduction of the activity at Rew-D with familiarity (Figure S2c). Our results suggest that, given more complex task structures, cognitive demands play an important role in determining the allocation of neural resources at reward locations.

### Hippocampal hierarchy changes with learning

We next asked how such local changes in neural activity affected the overall structure of the hippocampal code. We, in particular, focused on the relationship between the representations of different variables: were some representations subordinate, and thus nested, within the other representations that were dominant, and, if so, did this structure change with learning (Figure 1b-c). To formalize this question, we performed unsupervised hierarchical clustering on the neural activity PVs (Figure 3a-b) (20, 36). This method allowed us to organize PVs into clusters of growing size, from most similar to most different, in terms of population activity. We could then test whether task variables were drivers of this structure, by checking which variable values were shared between PVs of the same cluster.

**Figure 3.**
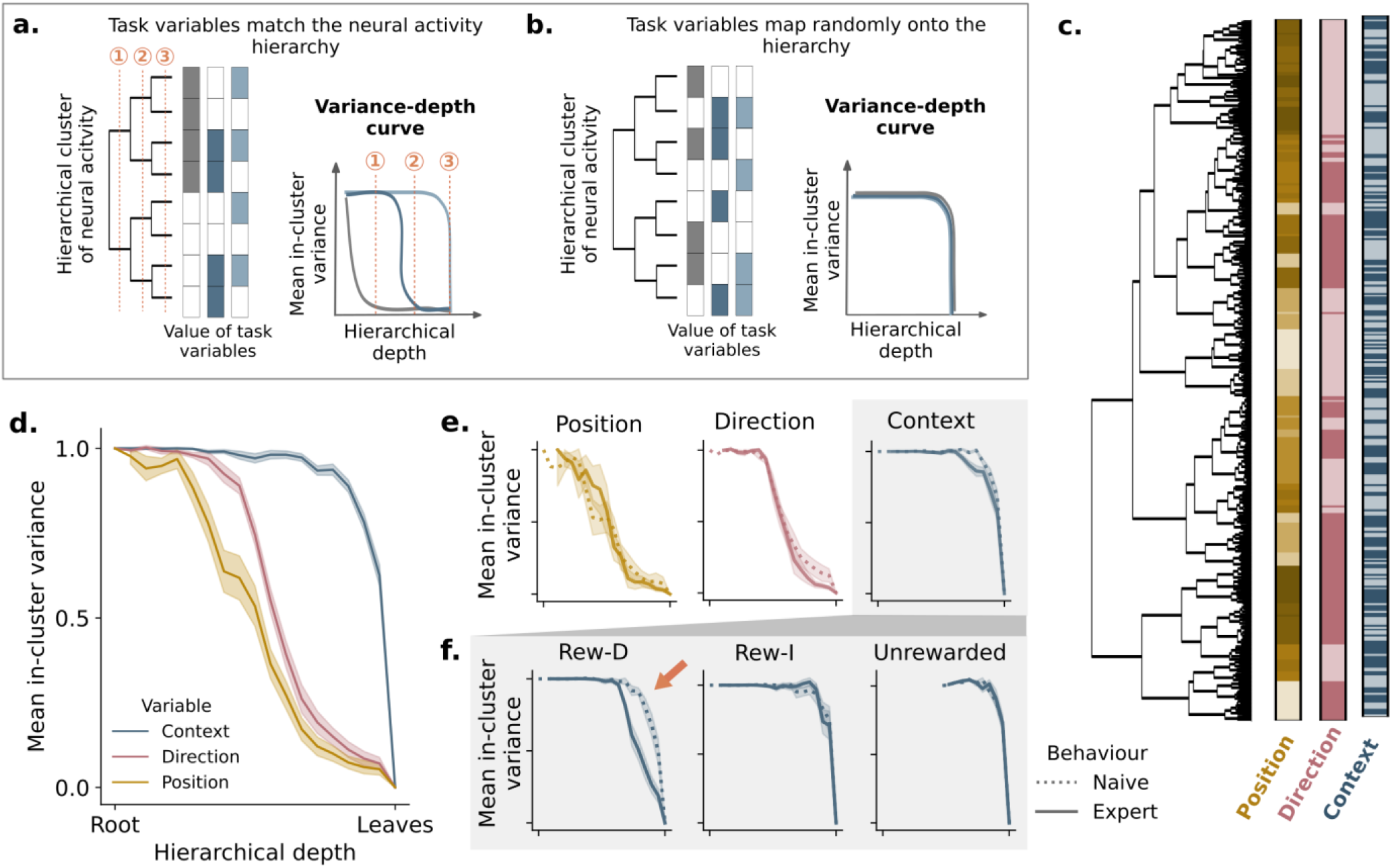
Changes in neural representation hierarchy with learning. **a-b)** Hierarchically-organized neural representation can be revealed by unsupervised clustering. **a)** Schematic of population vector clustering (dendrogram at left) with matching task variables (colored indicators at right; not used for clustering) that are perfectly hierarchically represented. Within-cluster variance of variable labels drops at defined hierarchical depth characteristic for each task variable. **b)** Same as (a) but for the case where there is no hierarchical representation; within-cluster variance drops at a similar, low-hierarchical level for all variables. **c)** Example dendrogram calculated from an Expert behavioral session, with the corresponding task variable labels. **d)** Label variance drops at characteristic levels of the hierarchy for different variables (color = mean over all sessions; shaded areas indicate SEM over all sessions in each behavior group, n=8 for each group). **e)** Same as (d), but separating out naive (dotted) and expert (solid) sessions. **f)** Variance of the “context” label for naive and expert sessions as in figure (e) but at branches that contain specific locations (shaded areas around the lines indicate SEM over all sessions in each behavior group, n=8 for each group). The representation of context at the context-dependent reward location (Rew-D) moves significantly towards higher hierarchical level (arrow).

We used PVs calculated for each trial separately to capture fluctuations occurring throughout the session. We also monitored the distribution of the PV-associated labels – context, direction, and position, which were not used for clustering – along the hierarchical tree. Position labels were coarse-grained to 9 possible values, corresponding to 40 cm bins. At each hierarchical level, we measured the variability of labels within each separate cluster. By definition, variance is maximal at the top level, where all population vectors form a single cluster, and zero at the lowest level, where each population vector forms an individual “cluster”. We refer to this relationship between variance and hierarchical depth as the variance-depth curve, a measure that allowed us to map task variables to the neural activity hierarchy. An early drop in variance with tree depth is an indication that a variable sits on top of the hierarchy, i.e., it drives the largest separation in neural activity (Figure 3a). Conversely, a later drop in variance can indicate that a variable sits at the bottom of the hierarchy (Figure 3a); it can also imply that the code maps randomly into the hierarchy, if that is the case for all variables (Figure 3b). If two variables drive equal levels of neural activity variance and/or are represented in largely disjoint areas of the activity space, then their label variance should follow similar curves (Figure 3b).

In Figure 3c, we show the neural activity hierarchy in an *expert* session. In this example, we can visualize a hierarchical distribution of task variables, which resembles Figure 3a; position and direction clustered activity within large branches of the tree, whilst contexts separated the activity closer to the leaves. This relationship was unaltered throughout the different learning stages in all animals, as quantified by the variance-depth curve (Figure 3d-e), indicating that position was the main external driver of hippocampal variability regardless of task learning. Within the representation of nearby positions, direction was the next driver of variability, and only at each position/direction did context have a detectable effect.

While the order of the variables remained constant within the hierarchy, changes in context encoding correlated with increasing performance at mid-hierarchy levels: there was an earlier drop in variance for context encoding in the *expert* quantile of trials (Figure 3e). This was not true for the other two variables, indicating that the effects of learning on the hierarchy were mostly restricted to the encoding of context (Figure 3e). Given our cosine similarity results shown previously (Figure 2f), we wondered whether the context grouped PVs earlier in the hierarchy, specifically at those positions where the cosine similarity between contexts decreased. This was indeed the case: branches that contained PVs from Rew-Ds showed lower context variance earlier in the hierarchy than those containing Rew-I position or no reward position PVs (Figure 3f). The same was also true when using PVs from the “speed-equalized” model, showing that differences in speed at those locations were not sufficient to explain the differences in firing between the two contexts (Figure S4). Taken together, this suggested that the hierarchy of the representations in the hippocampus could be altered locally via learning, according to the task demand, and at the same timescale at which behavioral performance changed.

### Changes in encoding hierarchy benefit downstream decoding

Cosine similarity between PVs (Figure 2f) and the representational hierarchy for context (Figure 3) changed over learning sessions, but these changes were not strong enough to prioritize context over position or direction. This could indicate that the hippocampus contributed to the context code but was not directly responsible for changes in task performance over days. Alternatively, albeit small, the changes in context representation were sufficient to support changes in behavior. To disentangle these hypotheses, we took the perspective of a downstream brain area, which has to use hippocampal information to trigger a behavioral response. A linearly decodable output is the simplest and arguably one of the most efficient codes that can plausibly be read out by downstream areas (43), so we used linear decoders – instantiated as Support Vector Machines (SVMs) – to decode behavior variables from the hippocampal population activity. We call these “global decoders” since they were trained on the entire data of a given session (Figure 4a, middle). Here, we used the “speed-equalized” PVs, to make sure we did not decode differences emerging from speed instead of the true context code. We trained the decoders using bootstrapped sampling and reported the classification accuracy in the cross-validation held-out test data, which were randomly chosen PVs not used for training the decoder. We first decoded each variable independently and found that all three variables (position, direction, and context) were decodable above chance at all stages of learning (Figure 4b). Notably, position and context were decoded at slightly worse accuracy in the speed-equalized case shown here compared to the original, non-speed-equalized data, indicating that differences in speed affected their linear separability (Figure S5b). We observed a trend for improved context decoding with better behavior performance (Figure 4c), but it was not significant with the “global decoder.” Furthermore, position and direction representations were stable already on the first day and were not directly related to performance gains over longer timescales (Figure 4c).

**Figure 4.**
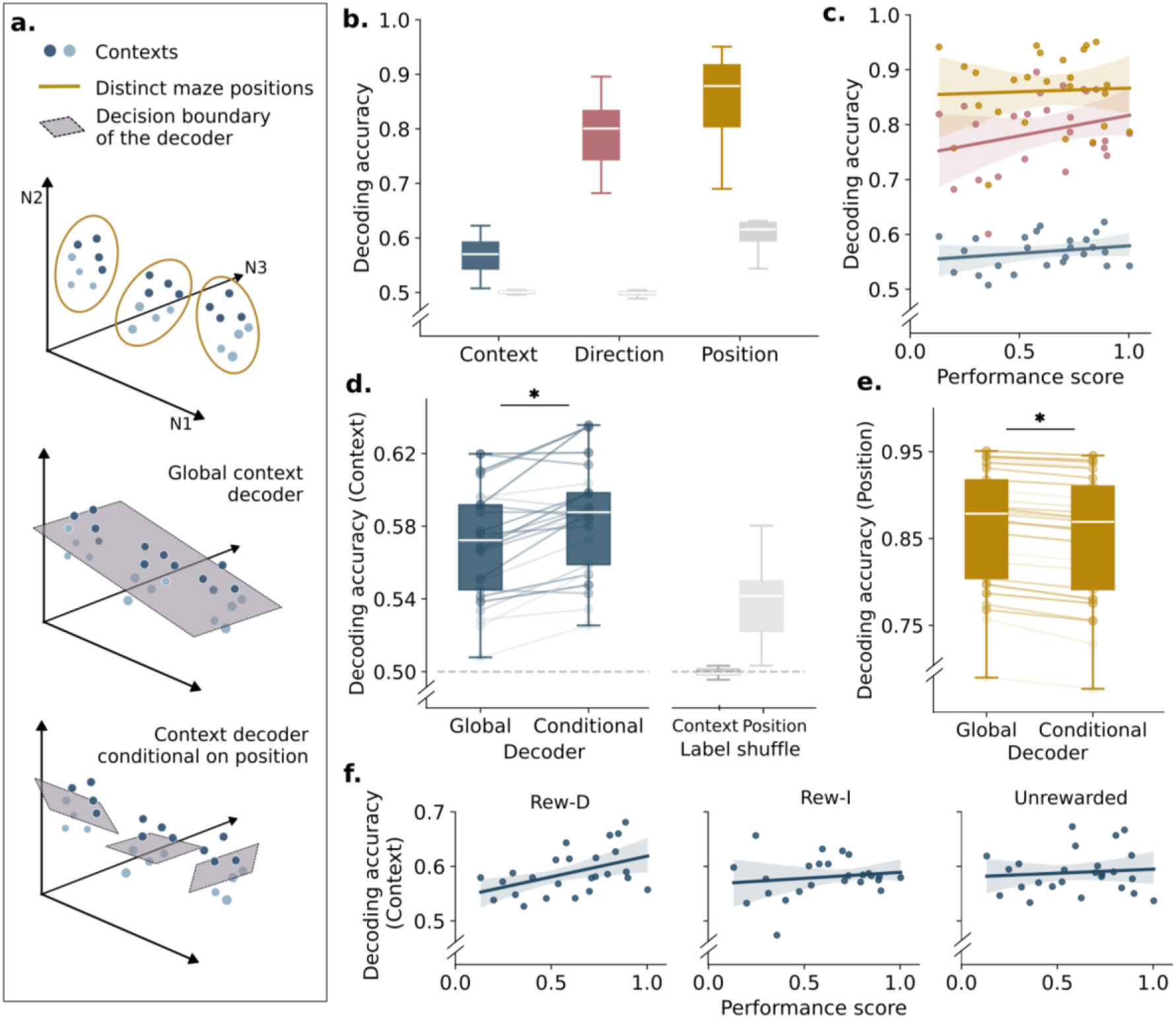
Global versus conditional decoding of task variables from neural activity. **a)** (Top) A possible hierarchical neural representation of position (yellow circles) and context (blue shade); each dot is a high-dimensional population vector depicted here in three dimensions. (Middle) A global linear decoder uses the same separating hyperplane (gray) to classify context across the entire set of population vectors. (Bottom) A conditional decoder consists of a set of linear decoders, each trained on a subset of data that shares the same label for the conditional variable (here, same position). Here we use cross-validated SVM classifiers with balancing for decoding either binary labels for context and direction, or 9 discrete labels for position (see Methods). **b)** Global decoder accuracy for each variable (context, direction, position) for all sessions (n=25) compared to decoding accuracy when the labels in the training set are randomly shuffled (gray boxes). Accuracy was evaluated over withheld (test) data as the mean absolute difference between decoded and real labels divided by maximum possible difference. All comparison to shuffle p<0.01, one-sided paired t-test with Bonferroni correction for multiple comparisons. **c)** Global decoder accuracy for each variable (context, direction, position) as a function of behavior performance. Correlations were n.s. for all three variables. **d)** Comparison of context decoding with global vs. conditional decoder (conditioning on position). The global decoder bar is the same as in panel (b). Line shading indicates behavior performance for each session (stronger colors = higher performance score). Gray bars indicate the conditional decoder performance when either context or position labels were shuffled. Global vs Conditional p < 3.80e^-5^, and all differences from shuffles p<<0.01, one-sided paired t-test with Bonferroni correction for multiple comparisons. **e)** Same as (d) but with position as the decoded variable and context as the conditional variable, showing that the conditional decoder actually does worse in this case. Global vs Conditional p<2.27e^-11^, one-sided paired t-test. **f)** Accuracy of context decoding conditional on position at specific reward positions as a function of the performance score. Rew-D R=0.70 p<9e^-4^; Rew-I n.s.; Unrewarded n.s.. All regression R and p-values were calculated with partial correlation, controlling for the number of pyramidal neurons recorded and animal IDs. Shaded areas indicate a 95% CI.

### Context information is local and task-specific

Given that the changes in the encoding hierarchy were detected mostly at specific maze locations – the Rew-Ds –, we wondered whether a decoding scheme that takes the encoding hierarchy into account could perform better than our global decoder. Specifically, we asked whether knowledge about the dominant variables, which were better decoded and lay higher in the hierarchy, could aid the decoding of context. To test this, we defined a “conditional decoder”: for each of the “decoded” variables we used before, multiple SVMs were trained, each on data that shared the same value of a “conditional” variable (Figure 4a, bottom). As an example, if we tried to decode context conditional on maze position, it means that, for each position, a separate SVM was trained on data coming from either of the two contexts. Although such a decoding model had more parameters, it had proportionally less data to train each SVM on, which could lead to low accuracy on the cross-validation data due to overfitting. Yet this was not what we observed: the conditional decoder outperformed the global decoder when predicting the context from held-out test data (Figure 4d), suggesting that reliably-decoded variables at the top of the hierarchy – such as position – could be used to facilitate decoding of other, lower variance variables. It is important to note that the opposite statement did not hold. Knowing context did not significantly improve the decoding of position or direction – actually, the corresponding conditional decoders performed worse on the held-out test set (Figures 4e and S5a).

The higher accuracy of the conditional decoder in comparison to the global case confirmed that context is encoded differently at each maze position. When we examined the performance of the decoder at individual positions, we observed better decoding of context at Rew-D, but not at Rew-I, on trials with high performance (Figure 4f). At other maze positions, we also detected a trend of improved context decoding with learning, but it was not statistically significant. Taken together, our results confirm that context influences neural activity in a position-specific and task-dependent manner. At Rew-I, for example, knowing context is not necessary for decision-making and, consequently, it does not need to be – and in fact it *is* not – better encoded, despite the reward being unconditionally present at this position.

The analyses performed so far quantified context decoding conditional on the *actual* position of the animal. In reality, a biologically plausible implementation would not have access to the *actual* position of the animal; instead, it would need to infer this position from the neural encoding first, in order to subsequently decode context, conditional on this putative, inferred position, later. When implemented in this two-step, sequential procedure that respects the encoding hierarchy, context decoding was highly efficient (Figure S5c). Interpretation-wise, such a sequential approach is only locally linear: while at each position, a linear context decoder is employed, taken together, context decoding is a globally non-linear operation. An alternative to such a sequential, locally-linear decoding procedure that respects the encoding hierarchy is a full joint decoder which linearly decoders a combined position+context label. Although such a decoder yielded similar results (Figure S5c), it requires the update of a larger number of parameters to adapt to the local changes in representation happening during learning (see Extended Methods), speaking in favor of a sequential approach.

### Hierarchical representation gets more efficient with learning

In our final analysis, we tried to elucidate the advantages of the hierarchical code structure and its changes with learning. When we trained context decoders conditionally on position, we provided each decoder with only a subset of PVs, all collected at the same position. Effectively, this means that in each such subset we removed the variance in neural activity that was driven by space modulation, allowing the conditional decoders to more easily extract context information from the same PVs. To better understand this decomposition, we performed an unsupervised dimensionality-reduction using Principal Component Analysis (PCA) to obtain directions of the highest neural activity variance, by making no use of task-related information in the process. We asked whether the identified directions of highest population activity variance contained information about task-relevant variables and how learning affected this correlation. To this end, we projected raw data onto the identified PCA directions and tested how our decoders performed when trained on differently-sized subspaces of the population activity.

An example projection of PVs for an entire session onto the first three principal components (PCs) is shown in Figures 5a-b. PVs corresponding to different maze positions were well separated along the top PC axes (Figure 5a). This is in accordance with our decoding results, where the position was the best decoded variable on any session. In the example we show, it was even possible to identify five different segments of the activity along the first three PCs, corresponding to the five segments of the real maze (other examples in Figure S6). Conversely, no clear separation of the neural activity from different contexts could be observed (Figure 5b).

**Figure 5.**
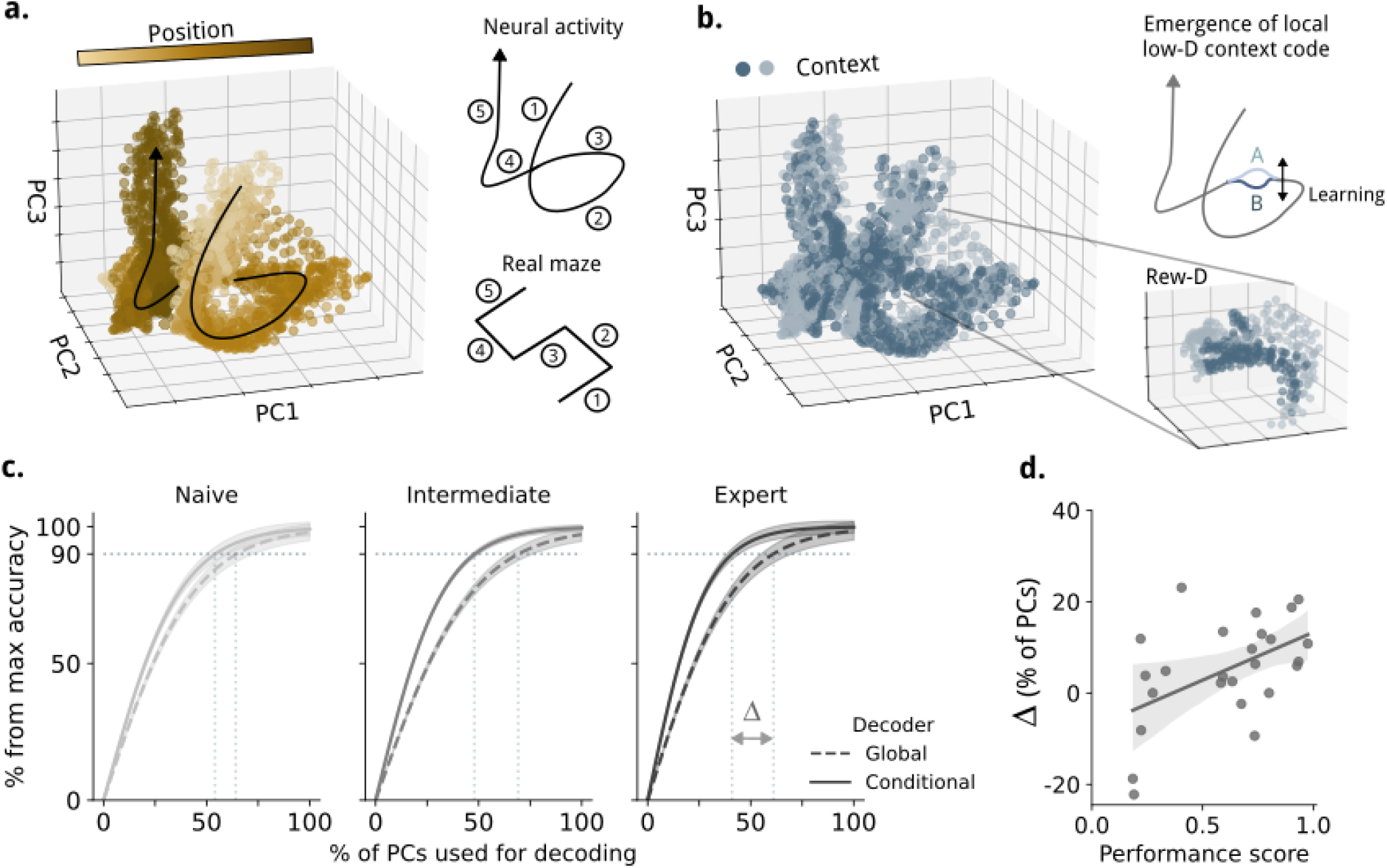
Decoding from dimensionally-reduced data. **a)** Example PCA projections for raw-data population vectors (PVs, calculated as average firing rates in each 4cm position bin in each individual trial) in one expert session; colors indicate true animal position. PC-projected neural activity can be partitioned into 5 segments corresponding to the real maze (diagram at right). **b)** Same as (a), but with data points colored by context instead of position. No global separation between contexts is visible in the first 3 PCs, but we hypothesized that a local low-dimensional separation could be possible, as we can see when we plot only the Rew-D location (zoom-in panel). **c)** Training increases fraction of neural activity variance devoted to context coding in a position-dependent manner. Global (dashed) and conditional (solid) context decoder cross-validated accuracy for naive (n=8 sessions), intermediate (n=9), and expert (n=8) sessions (left to right), shown as a function of the fraction of PCs used for decoding and normalized to accuracy achievable on the complete dataset by each specific decoder (using all PCs). Shaded areas around the lines indicate SEM over all sessions in each behavior group. Vertical dotted lines show the fraction of PCs needed by the global and conditional context decoders to reach 90% of their respective maximum accuracy. **d)** Difference Δ between the two decoders in terms of required number of PCs at 90% of maximum accuracy. R=0.41 p<0.05. Shaded areas = 95% regression CI. The regression lines result from a partial regression controlling for the number or pyramidal cells recorded in each session.

When we zoomed into the population vectors extracted from a single maze position, we observed that, for some positions, there indeed was a separation of the remaining task variables in the first PCs (Figure S6). If this separation was consistent but not aligned between different positions, a conditional decoder would be expected to outperform the global decoder as reported previously – but conditional decoder should *also* outperform the global one when trained only on a small subset of top PCs and not only when trained on the entire dataset. To test this hypothesis we measured how many PCs were necessary for each decoder to reach 90% of its maximum performance (Figure 5c). As expected, the conditional decoder required fewer PCs to reach 90% accuracy for decoding context, when compared to the global decoder. This is not surprising given that position, which was always highly decodable, was already well separated along the first principal components (Figure S5d). Our result also implies that context was also partially separated within this same subspace, but the context separation hyperplane was not necessarily aligned across positions. As a result, a globally linear separation of context (global decoder) within the first PCs was suboptimal compared to the locally linear one (conditional decoder). More interestingly, as behavioral performance increased, the difference between the two decoders in the number of PCs required to reach this 90% threshold substantially increased (Figure 5d). This suggests that spatially-modulated cells started carrying more contextual information as learning progressed. At the population level, this indicates that variables were represented – at least in part – within the same linear subspaces of activity and in a hierarchy, and that learning increased the overlap between the representations of associated variables.

## Discussion

In this paper we showed how population-level representations in hippocampal principal cells reorganize as the learning progresses over multiple days. Using a space-context association task, we observed a robust hierarchy in the representation of behaviorally relevant variables: physical position was the largest driver of separation between representations; within the representation for each given position, we found a subordinate separation between representations for the two different movement directions, and, finally, a lowest-order separation between the two task-relevant contexts. Successful learning did not change this overall organization of the neural code. Yet, as the performance improved, context representations did separate substantially at task-relevant positions, at the deeper levels of the neural activity hierarchy. At these task-relevant positions, the representation of context migrated towards the axes of higher neural activity variance. We demonstrated that this type of hierarchical representation and its reorganization facilitates downstream decoding: as the learning progressed, downstream areas could still use the same circuits to decode position and direction, and only context decoding needed to be refined using readily available information about the other variables. Together, these results show that the hippocampal code adapts to task requirements occurring at the timescale of behavioral changes; this is implemented through local refinements that are commensurate with the improvements in behavior performance.

The cognitive benefits of hierarchical representation structures have been described in multiple studies in humans and other animals (20, 36, 44–46): they allow prioritization of variables according to their relevance for behavior. In our case, we suggest that the representational hierarchy in the hippocampus prioritizes essential behaviors (e.g., position awareness); nevertheless, it is simultaneously adaptable to trained behaviors. We have shown that position and movement direction were at higher hierarchical levels at all learning stages, which is not surprising given the relevance of these variables to most rat behaviors. Nonetheless, learning led to selective changes in the representation of context, initially encoded at the bottom of the hierarchy. Context drove increasingly more neural variability at locations where it was consequential for reward, and this adaptation matched the changes in behavior performance. Taken together, this indicates that the hippocampus can accommodate behavioral demands within its representation hierarchy.

In contrast to our results, Mckenzie et al. (20) previously observed that context representation is at the top of the hierarchy in well-trained animals. The difference between the two studies is likely due to different task design: McKenzie et al. defined different contexts by distinct physical environments, and learning involved differentiating the valence of different objects within each of those environments. Therefore, from the beginning, distinct cognitive maps were assigned to different contexts, while the learning of valence-object associations resulted in local changes in the representations at the bottom of the hierarchy. However, in our case, the same physical maze was used and context was defined by different floor cues. Often, changes in sensory cues in an identical environment trigger rate remapping (27). Rate remapping does not change place field location, it only alters the firing rate of place cells, which may explain why context was represented at the bottom of the hierarchy. Although context was represented at different hierarchical levels in our and the McKenzie et al. studies, in both studies the result of associative learning (object vs. valence in the McKenzie case and context vs. location here) was a change of encoding at the bottom of the hierarchy.

These and past results suggest that the hippocampus can reorganize representations as a result of learning in two distinct ways. In cases when the animal detects a novel environment or an entirely novel learning situation in a familiar environment, a novel representation is formed rapidly. However, once this representation is formed, slow associative learning triggers only localized refinement of the existing representation. This small and local adaptation of the code is reminiscent of the “lazy” learning regime observed in the training of artificial networks, where high classification accuracy is achieved by small changes in the network internal representation (47). However, lazy regimes in artificial networks are associated with fast representation learning and therefore do not fully explain our observations. Moreover, the learning regime can be different in other brain areas: in the human and monkey prefrontal cortex, contextual learning resulted in larger representation changes, resembling instead a “rich” learning regime from artificial neural networks trained on a similar task (48).

Our work also suggests that map refinement during learning is accompanied by dimensionality reduction in the representation of context. Recent works have shown that, while behavior sets an upper bound on the dimensionality of neural activity (49), the actual dimensionality of associative brain areas balances compression (fewer dimensions) and representational flexibility (more dimensions) to represent complex tasks (35). This usually results in more than one dimension representing single task variables, which we also observed in our data, albeit only as measured in the PCA space. Moreover, a reduction in dimensionality with learning has been observed in multiple brain areas (35, 50, 51) and in artificial neural networks (for a review, see (15)). It has been proposed that this can reduce the resources necessary to perform learned tasks and improve generalization (35). Such dimensionality and consequently redundancy reduction can also be understood in terms of an efficient code where optimization is not exclusively based on input statistics, but actually takes the task goal into account (52, 53). This could explain the selective activity gain increase restricted only to Rew-D as opposed to Rew-I positions when optimizing for this specific task. Although our results do not preclude alternative explanations, they are consistent with these previous theories.

## Materials and Methods

### Animals

All experiments were conducted on 5 Wild-type, Long Evans Rats of 9 - 13 weeks of age and weighing 300-350g. Animals were caged with littermates before implantation surgery and isolated afterwards. They were kept in a dedicated animal room under a 12 hour light/dark cycle. All animal procedures were carried out in accordance with the Austrian federal law for experiments with live animals, under the license number BMBWF-66.018/0018-V/3b/2019.

### Behavior apparatus

The maze consisted of a raised S-shaped platform, 10 cm wide and 360 cm long. The shape was formed by 5 arms of 80 cm, each overlapping 10cm at the corners. The track was surrounded by 30 cm-high transparent walls and a square box on either end, each 20 cm wide surrounded by high walls and a door to the maze. Each maze arm contained 7 wells 10cm apart (15 cm from the start box). A photo of the maze is shown in Figure S1a. Wells were 5cm wide and 5cm deep and were filled with sand. Extra tiles could be added to the maze surface, changing its texture and color and covering subsets of the wells. In this experiment, the possible tiles (determining context) were two: a black, hard plastic tile with equidistant grooves of around 1mm; and a white, EVA-foam sheet tile with a smooth surface. Fixed distal cues were hung around the room to facilitate distinguishing the overall position of the maze.

### Habituation

Before the start of training, animals were habituated to a blank version of the maze, where there were no contextual tiles. In this period, they learned to dig wells of sand in the home cage and in the maze. They were exposed to wells with progressively less food/sand ratio until only one flake was present per well. The maze wells available during habituation were not available during training. In this period, they were trained to run towards the end box after finding food. No association task was introduced during habituation.

### Behavior task

During habituation animals were put under a food restriction protocol, and once animals reached 85-90% of their pre-surgery weight, we started the training and recordings. We use the term “behavior session” to refer to a day of training encompassing 40 trials, in alternating blocks of 5 trials for each context. In order for the animals to learn this task appropriately, food was laid on top of the sand for the first 10 trials of the first training day and then buried halfway deep for the remainder of the experiment. The starting position (left/right box) was random but there were an equal number of trials from each.

On a single trial, animals had to use the color and texture of the maze as cues to decide where one of the rewards would be located (context-dependent). A second reward would be found at a fixed location, independent of the cues (context-independent). A total of 8 sand-filled wells were available at each context, 4 of which overlapped between contexts in terms of position (the three rewarded wells plus a control well). Animals were only allowed to dig two wells in a trial; after the second choice was made all other wells were covered and made unavailable. Nevertheless, animals were allowed to go back and forth on the track and dig the chosen wells as many times as they wanted, for up to 5 minutes. Only one corn flake was made available at each reward well. Animals terminated a trial by running to the box at the opposite end from which they started, where they would find a sucrose pellet if the two wells dug were correct. In case of an incorrect choice, there was no reward at the end box. The door to the end box was open only once the animal made two choices. If the animal did not complete the task in 5 minutes, the trial was terminated and no further reward was provided in the trial. We made sure that no residual odors were guiding the animal’s choice, by mixing the sand between wells and cleaning the maze regularly.

### Performance score and grouping

To assess how many trials an animal performed using a contextual strategy, we defined a performance score which accounts for the fact that a certain number of correct trials could be achieved without using context for decision-making. In naive sessions animals mostly employed what we call an “eager” strategy, in which they dug the first 2 out of the three possible reward wells, disconsidering the current contextual cues. A “contextual” strategy would involve digging the context-independent reward and the context-dependent reward corresponding to the current sensory cues, regardless of the order they appear, given the current running direction. We call trial categories (defined by direction and context) unambiguous if these two strategies would yield different digging choices in that trial and ambiguous when both would yield the same choice outcome. The ambiguity refers to the possibility of distinguishing the strategies, but not the actual behavior of the animal.

For each session, we then calculated the proportion of trials in which the behavior actually corresponded to each strategy. Note that the proportion of correct trials is synonymous with the proportion of “context” strategy trials. From the unambiguous categories, we calculated the proportion of trials which were eager trials. Note that these trials were incorrect, but other types of errors were also possible such as digging only one or no well. These other errors were not evidence for either strategy, but still accounted for the total number of trials. We then assumed this same proportion of eager trials was true in the ambiguous categories. So the final score was calculated as:

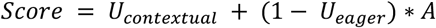

where *U*_*s*_ indicates the proportion of unambiguous trials where the behavior corresponded to strategy *s*, and *A* the proportion of trials in the ambiguous trial categories which the animal made a correct choice, and therefore were consistent with both strategies. Lastly, we separated the sessions (N=25) into three quantiles of performance score, *naive* (N=8 sessions, from 5 animals), *intermediate* (N=9 sessions, from 5 animals) and *expert* (N=8 sessions, from 4 animals).

### Behavior measures

For the mean speed calculation per session we first filtered out bins where the animal was at the start/end boxes, as well as when the animal was immobile (< 3cm/s) or not running in the main direction of running of a given trial (e.g. running towards the left in a trial that started on the left). We calculated the average immediate speed over all remaining bins.

For the calculation of time spent at rewards we summed all immobile periods spent in a radius of 10 cm from the reward locations the animal dug in a given trial. The remaining time spent in the maze we denote trial time (excluding reward).

### Microdrive implantation surgery

We implanted 16-tetrode microdrives in the dorsal CA1 area of the hippocampus of the rats. Two animals also received a second set of 16-tetrodes implanted to the retrosplenial cortex. The surgery protocol implemented has been described in previous work (54, 55). In short, animals were put under isoflurane anesthesia and received rehydration with a saline/glucose solution mix every 2 hours during the surgery. Analgesia was achieved with metamizol and buprenorphine. Medication dosage was calculated according to the animal’s body weight based on the Austrian animal law. With the help of a stereotaxic device and a dental drill, the craniotomies were made over the hippocampus at AP: -2.5 to -4.5 and ML: 1.0 to 4.0 from Bregma. During the surgery tetrodes were lowered 1.0 to 1.3 mm into the brain; the remaining depth to reach the CA1 stratum pyramidale was achieved by tuning tetrode depth on the days following surgery. After surgery, animals were allowed to recover for at least 7 days before the start of behavior training.

### Electrophysiological recordings and position tracking

In vivo extracellular electrophysiological recordings were made using an Axona Ltd. recording apparatus. We used a tethered system which included a pre-amplification of the analogue signal directly at the animal’s head, then a second step of amplification and digitization of the signal before relaying it to the computer. The neural signal was recorded at a 24kHz rate. For position tracking, the animal’s headstage contained a pair of LEDs, one on each side. The LEDs were of different sizes, to allow identification of the left and right sides of the animal. Using an overhead camera and low light conditions, the system captured the position of the LEDs throughout the task, at a 50Hz rate.

### Spike extraction and spike sorting

Spike extraction and automated spike sorting were performed using MountainSort software (56). We also performed manual curation of the electrophysiological units, filtering units and classifying into putative pyramidal or interneuron based on spike waveshape, firing autocorrelogram and firing rate as previously described (57).

### GLM rate maps

We utilized a GLM model to describe each cell’s firing probability as a function of trial, position, and speed. We excluded from the calculation bins when the animal ran in the opposite direction than that stipulated by the start to end box direction. We will describe in detail here the covariates used to fit the model and the parameters used, and refer the reader to other references for the details regarding the use, fitting, and theoretical background of GLMs (38, 39). The model describes, separately for each cell, the Poisson probability of observing *n* spikes in a 20ms time window, *τ* = 0.02 *s*, given covariates *θ*_*t*_

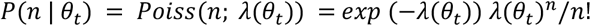

where

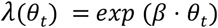

*β* represents the model coefficients, found by maximum likelihood (more on this below) and *θ*_*t*_ represents the covariates at time *t*, which are:

- Joint trial-position (one-hot variable): we binned the linearized position into 90 bins of 4 cm each. We allowed each unit to have a different parameter for each position on each of the 40 trials separately. This resulted in a 90*40-dim vector.
- Speed (one hot variable): we binned the speed, which was measured from the behavioral recording in 20ms time bins, in 10 non-overlapping and equally populated speed bins, excluding speeds below 3 cm/s.
- Constant: to capture any baseline activity.

The number of parameters of such models was therefore fixed to 90*40+10+1=3611. We utilized the routine GLM.fit_regularized offered by the package statsmodels v0.14 in Python 3.11 (58). We fitted the models by using an L2 regularization with parameter fixed to 10^-5^ for all but the constant covariate, which was not regularized. The regularization parameter was fixed by cross-validation, and selected so as to maximize the log-likelihood on the validation set with a grid search on the possible parameters (10^−8^, … 10^−3^).

### Population Vector (PV) calculation

From raw spiking data: the entire maze (360 cm) was divided into 90 spatial bins of 4cm each. For each trial and position, we calculated the mean firing rate of each neuron; the activity of all neurons formed a population vector of length *n*, where *n* is the number of putative pyramidal neurons in a given session. Each PV was assigned a set of labels: trial context, trial direction, and 2 position labels. The direction refers to the binary 1D directionality within the maze (towards the left or right end of the maze), which differs from the head direction in the real S-shaped environment; we excluded from the calculation periods when the animal ran in the opposite direction than that stipulated by the start to end box direction. The position labels were either the fine-grained (90 bins) or coarse grained values (9 bins). We indicate in the main text which of the two were used in each analysis.

“Speed-equalized” from the GLM: given the one-hot encoding, our GLM model yielded a coefficient for each position in each trial, as well as separate coefficients for each speed bin. The PVs from the GLM were calculated as:

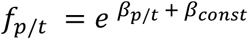

where *f*_*p*/*t*_ is the GLM estimated mean firing rate at position *p* at trial *t* and *β* are the model coefficients. This calculation corresponds to the mean firing at each bin excluding the estimated speed influence. The same number of PVs and labels were used as in the raw spiking data.

### Cosine similarity

Cosine similarity of the PVs was calculated creating a matrix of dimensions *n* x *p* for each trial category, where *n* was the number of neurons, *p* the number of position bins.

Each row of this matrix contained the mean rate map of each neuron in that category (4cm spatial bins, 90 bins in total). Cosine similarity was calculated between the matrix of categories of the same direction but different context or vice-versa. For each position *p*, we calculated the cosine similarity between the PV for that position and then averaged the similarity over all positions, obtaining the average similarity between trial categories. Shuffling of cell IDs was performed as a control. For comparison of running directions, the best similarity was taken from all off-diagonals to compensate for the place field shift (mean 34 ± 13cm). For more information on place field shift, see Lee et al. (59).

### Global and conditional decoding

We used Support Vector Machines (SVM) to decode task variables (i.e. coarse-grained position, context and direction) labels from the PVs calculated per position (90 x 4cm bins) and trial (40 trials) or their projections in a subset of the Principal Components. An SVM finds the best high-dimensional plane that separates the different categories, by finding the plane with the largest distance to points from any category (60). Over 100 repetitions, we randomly split the dataset in 70-30% to use as training and testing sets, respectively, making sure labels remained balanced in both subsets. Using a bootstrapping method, each subset was resampled in a stratified manner (the sampling keeps the label balance the same as in the original data). The same data splits and bootstrapped samples were used for training/testing the global and conditional decoder as well as for the shuffled label condition. The two decoding strategies employed were the following:

1. Global decoder: a standard SVM which was trained directly on the labels of the variable of interest to be decoded, from the entire set of training PVs in a behavior session.
2. Conditional decoder: each vector was assigned two labels, one for the conditional variable and one for the variable of interest. We subdivided the vectors into groups determined by their conditional variable. For each group, we trained a separate SVM. The accuracy was calculated as the average accuracy over all groups.

The decoding accuracy was calculated as the difference between the predicted labels and the real labels of each vector in the test set, divided by the maximum possible difference. For binary labels (context and direction), this difference was either 0 (match) or 1 (mismatch). In the shuffle condition, we calculated the average accuracy of each decoder when we randomly shuffled the labels for the variable of interest or the conditional variable. To control for the different number of neurons recorded in each session, all statistics that compare between sessions take this number as a control factor, as specified in the corresponding figure legends.

### Hierarchical clustering of population vectors

For the hierarchical clustering, we have used the PVs calculated as described above, calculated per trial per 40cm position bin (resulting in 40 trial x 9 bins = 360 PVs). The clustering algorithm did the following: Initially, each point represents a cluster of size 1 each; clusters are merged recursively following the Ward variance minimization criterion. In detail, denote a cluster as *X* = { *x*_1_, …, *x*_*k*_}, and compute its mean 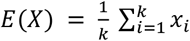 and its variance 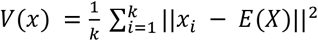, where || *x*_i_ − *E*(*X*)|| is the Euclidean distance between each point and the cluster center. At each step, the algorithm computes the variance of each possible merged pair of clusters *V*(*X*_*i*_ ∪ *X*_*j*_), and selects the pair to merge that yields the lowest increase in total variance. This procedure is computationally demanding; a simplified algorithm with linear complexity in the number of points has been proposed by Müllner (61) and is implemented in the Scipy library version 1.11 in the function cluster.hierarchy.linkage which was used here for analysis. This process is iterated over until all points are clustered together, and that is the root of the tree.

### Variance-depth curve

The hierarchical clustering procedure provided us with progressively finer subdivisions of PVs. We measured whether PVs within the same cluster, across depths, had homogeneous category labels. To do so, given a full hierarchical tree of PVs, we calculated the variance *W* of each label category *L* (position, direction, context) within each cluster at different tree depths. Note that the variance *V* used for hierarchical clustering refers to the variance in firing rates in each PV, whilst *W* refers to the variance of the label values. As one progresses from root to leaves, the number of separate clusters in the tree grows. At each depth we calculated the variance of the labels over all the PVs in each cluster separately and then averaged this value over all clusters, as follows:

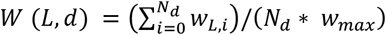

where *L* is the label category, *d* is the depth in the hierarchical tree, *N*_*d*_ is the number of clusters at that depth, *w*_*L*,*i*_ is the variance of the label *L* in each cluster *i* and *w*_*max*_ is the variance at the root. Each label *L* can take multiple values *l*. Direction and context are binary variables (*l* ∈ {0,1}) and position can take 9 discrete values (*l* ∈ {0,1, …,9}). We repeated this procedure at each depth (each new split of tree branches) until we reached the leaves, where variance is zero (only one PV per cluster).

### Statistics

The statistical tests used in each analysis are indicated in the corresponding figure legends.

### Code

All analysis were performed using custom made python scripts and the relevant code will be made available at: github.com/hchiossi/hpc-hierarchy

## Acknowledgments

We would like to thank Rebecca Morse for performing the recordings in one of the animals under the supervision of H.S.C.C., Jago Wallenschus for the technical support, especially with maze design, Wiktor Mlynarski for the advice and discussions and Andrea Cumpelik for suggestions during the writing. M.N. was supported by the Howard Hughes Medical Institute.

## Author Contributions

J.L.C. and G.T. secured funding. H.S.C.C. and J.L.C. designed the experiments. H.S.C.C. performed the experiments and data analysis. M.N. wrote the GLM model. M.N., J.L.C. and G.T. provided support to the analysis. H.S.C.C., M.N., J.L.C. and G.T. wrote the manuscript.

