## Supplementary Methods and Figures for "Learning reshapes the hippocampal representation hierarchy"

### Extended Methods

Here we explain the methods that only appear in the supporting figures.

*Successful refrain trials*

A behavior was labeled "refrain" when animals successfully held back from eagerly digging the incorrect reward well. More specifically, we measured the proportion of correct trials from all trials in which the incorrect contextual reward well was positioned before the correct one, given the trial running direction.

*Firing rate gain*

The average firing rate gain per position in each trial category was defined as the mean of the normalized rate map over all pyramidal neurons. If we denote $r_i$ the rate map vector for each neuron $i$, $\bar{r_i}$ its average, and $n$ the number of pyramidal neurons, the gain was calculated as follows:

$$gain \ = \ (\sum_i r_i/\overline{r_i})/n \text{ -1}$$

*Sequential decoding and decoding from joint labels*

In our analysis using conditional decoders, we used the real labels of the conditional variable, but in reality this variable also needs to be inferred by the network. We therefore created a sequential decoder, where we first used a global decoder for the conditional variable and then used the resulting predictions to train the conditional decoder as before. We performed this calculation on the real, raw data, not the speed-equalized GLM, to show that the neural activity as observed by the brain can be decoded in this manner. The first decoder was a global decoder trained using 5-fold cross validation, so that for each data point a predicted value was obtained from one of the folds.

For the global decoding using joint labels we created new labels which were a combination of the position and context of each PV. In this case $p$ x $c$ categories had to be decoded, where $p$ is the total number of positions and $c$ the number of contexts. This decoder was trained the same way as the global decoder described in the main Methods. In short, we trained a Support Vector Machine to decode $pc$ labels from the PVs, using 100 times bootstrapping samples and 70-30% train-test split for cross validation.

The joint decoder also benefits from the position-specificity of the context code, since these two variables become a single combined variable in this case. However, an important distinction has to be made between the conditional decoder and global decoder of joint variables. Multi-class SVM decoders were trained in a one-vs-all manner, which means that each variable requires $n-1$ decoders to be classified, where $n$ is the number of discrete values this variable can take. If we call $p$ the number of positions, $c$ the number of contexts and $d$ the number of movement directions, then in the global decoding of joint labels the total number of decoders necessary is $p\ d\ c\ -1$. In the conditional decoding, assuming decoding in the hierarchical order, the number of decoders necessary is the same$(p-1)\ +\ p\ (d-1)\ +\ pd\ (c\ -\ 1)$, which also results in $pdc\ -1$ decoders. However, they differ in terms of update. For example, if the context code changes in a single location, all $pdc-1$ global decoders need to be updated. In the conditional case, on the other hand, only the context decoders for that single position need to be updated, so $dc-1$ decoders. Therefore, conditional decoders require less changes in decoder parameters during learning, but achieve similar levels of decoding accuracy (Figure S5c).

### Supporting Figures

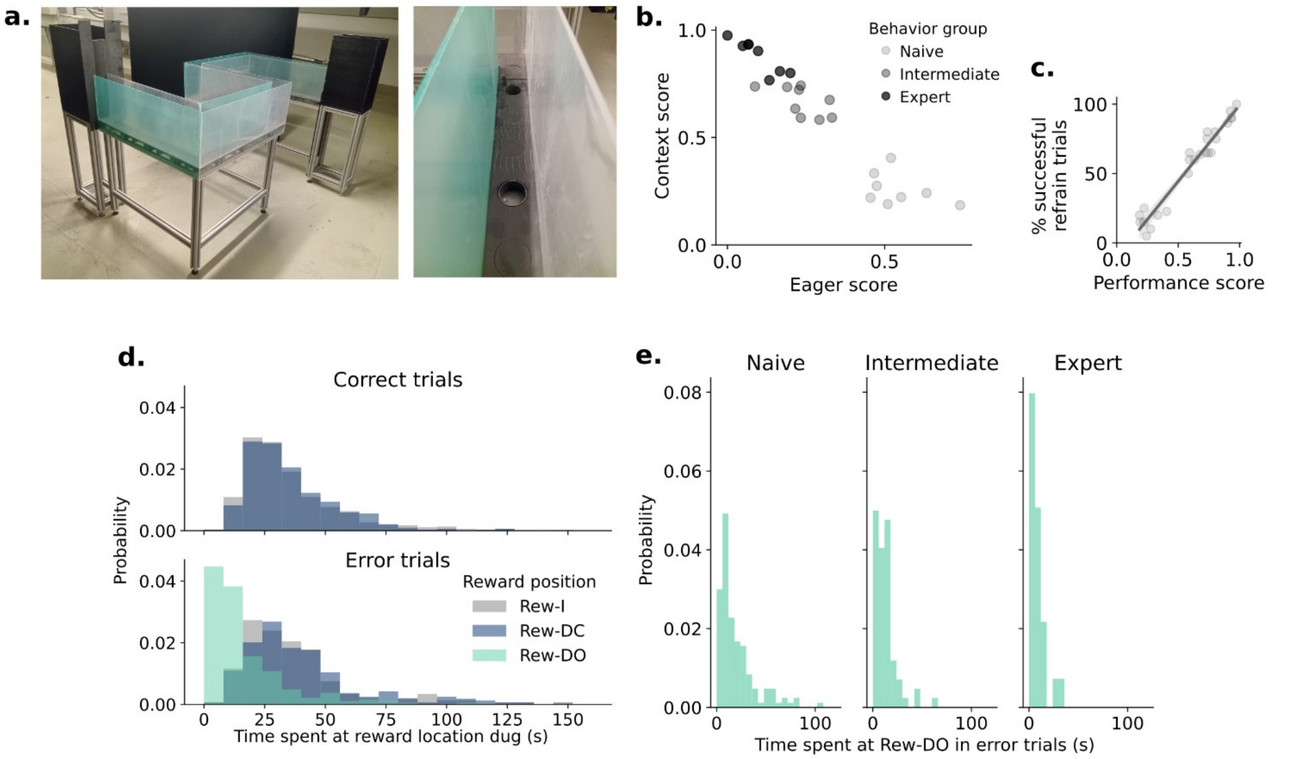


**Figure S1. Extended behavior measures. a)** Photo of the S-Maze and zoom-in of the track showing the empty wells where animals could dig. During the experiment the maze track was covered with different textures and the wells were filled with sand. **b)** Proportion of trials corresponding to a Contextual strategy (Context score) versus the proportion corresponding to an Eager strategy (Eager score) for each behavior session (N=25). The shades of gray indicate the different behavior quantiles to which these sessions were assigned. **c)** Refrain trials were those in which the incorrect contextual reward well was positioned before the correct one. The percentage of those trials in which the animal successfully refrained from digging the incorrect well strongly correlates with the animal's performance. $r=0.98$ $p=6e^{-14}$ (partial correlation controlling for animal ID). **d)** In correct trials (top) animals spent equal amounts of time around the Rew-I and Rew-DC (context-dependent reward of the Current context). In error trials (bottom), animals also spent time at the incorrect Rew-DO (context-dependent reward of the Opposite context), but less than in the other two reward locations, indicating that animals were likely aware of their mistakes. **e)** The time spent at Rew-DO during error decreased with improved behavior performance.

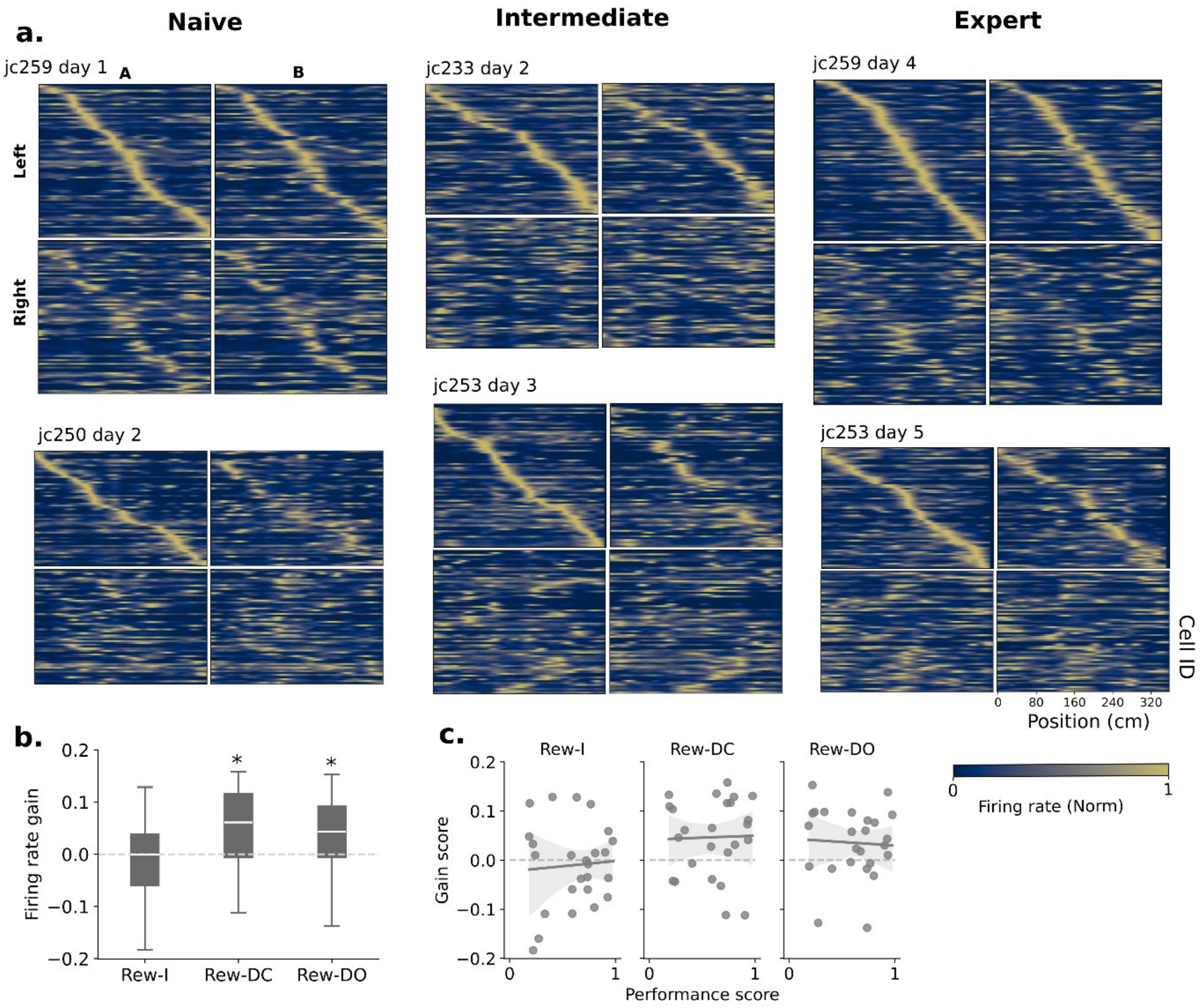


**Figure S2. Place cells and firing rate gain. a)** Examples of place cell population from different animals on different sessions. Cells are ordered according to the position of their firing rate peak at the Left/A trial category. **b)** Firing rate gain at the different reward locations, showing that at the contextual rewards firing was above average (of the current and opposite trial categories, Rew-DC and Rew-DO). A gain of 0 indicates the firing is equal to the mean firing on the entire maze, showing that there was no excessive firing at Rew-I, despite the fact that animals found food there. Rew-I n.s. Rew-DC $p<0.01$, Rew-DO $p<0.02$, one-sided t-test. **c)** There was no significant change in the gain at reward locations between sessions with different performance scores. Partial regressions controlling for the factor animal, all results are n.s..

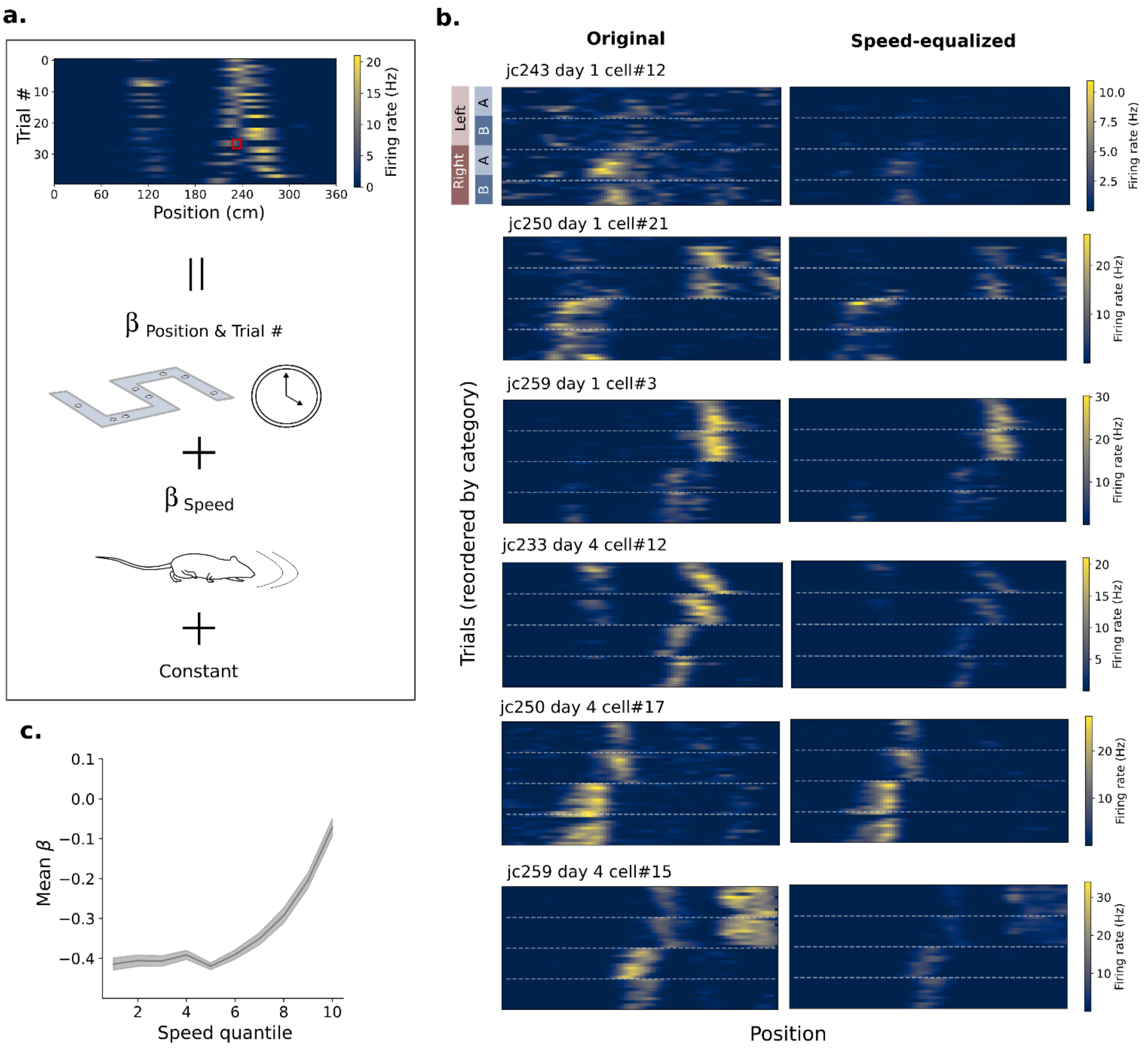


**Figure S3. GLM model for obtaining speed-equalized rate maps. a)** Schematic of the GLM model. The model to describe each cell's firing probability as a function of trial & position (as a single variable), and speed (see Methods). **b)** Example cells comparing the rate maps per position and trial obtained by using the raw spike counts and those obtained by the GLM. By removing the contribution of any speed, there was generally an attenuation of the peak firing rate, which varied between different trials & positions, given their difference in original speed. The strength of this attenuation also differed between cells. See Methods for details on the implementation. **c)** The GLM coefficient for each speed bin (10 equally populated quantiles) increases for larger speeds, indicating that higher speeds led to higher firing rates. Shaded areas indicate SEM (N=25 sessions).

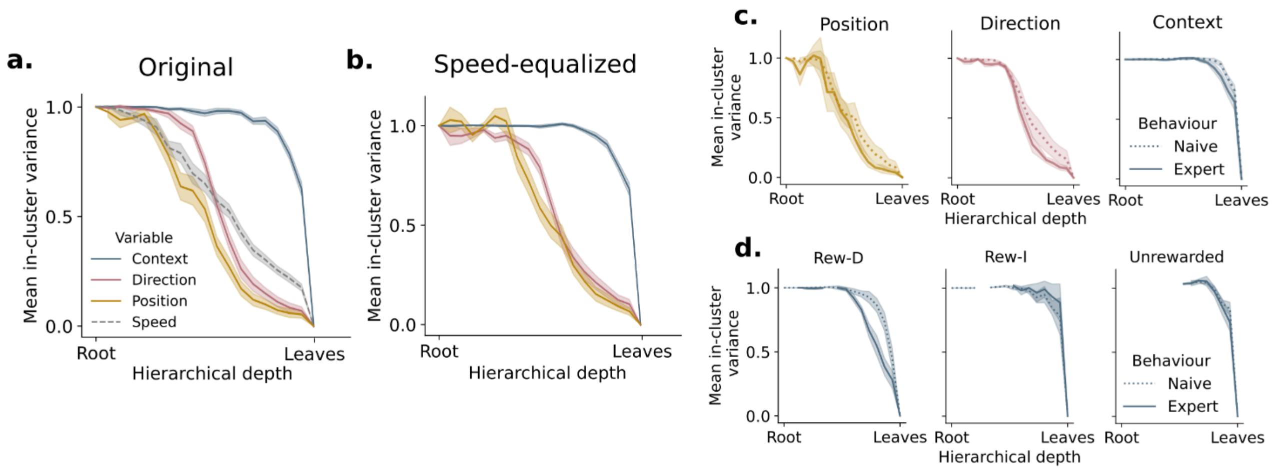


**Figure S4. Population vector hierarchy on GLM data. a)** Same plot as in the Main Figure 3d, but also showing the variance-depth curve for the Speed variable (10 bins). **b)** Variance-depth curve for the speed-equalized PVs calculated from the GLM. The right shift in the position curve compared to (a) indicates that the difference between positions in the original data can be partially explained by the differences in speed at each part of the maze. **c)** Same as main figure 3e, but calculated for the speed-equalized rate maps. The changes between naive and expert behaviors in the speed-equalized PVs remains very similar to that in the original data. **d)** Same as main figure 3f, but calculated for the speed equalized rate maps. This shows that the difference in the hierarchy at the Rew-D is maintained in the speed-equalized model, showing that differences in speed at those locations are not sufficient to explain the differences in firing between the two contexts.

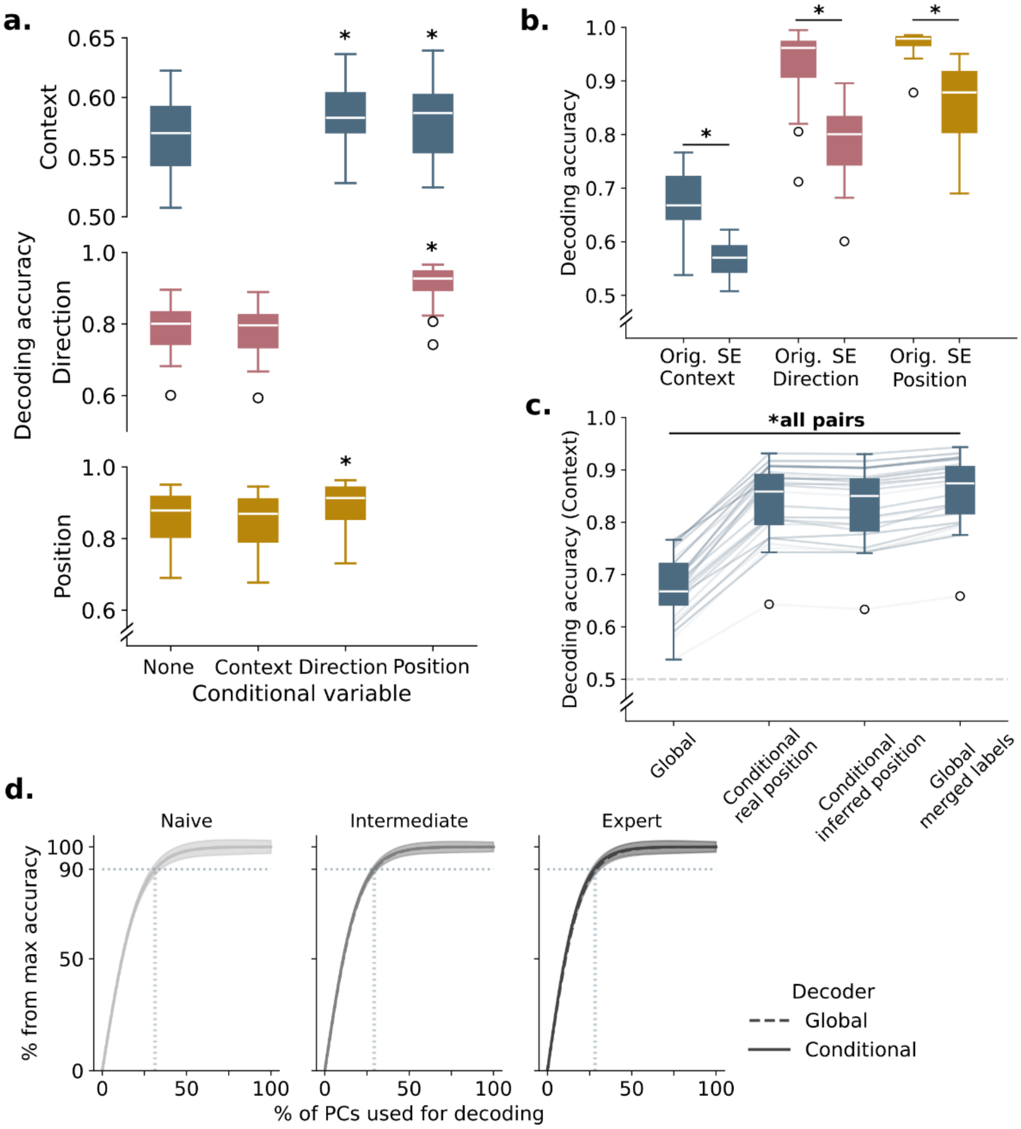


**Figure S5. Extended decoding results. a)** Decoding in the hierarchical order increases decoding accuracy. Context is better decoded conditional both on Side or Position, but the opposite is not true. Here "None" for the conditional variable refers to the Global decoder. All comparisons to Global decoder ('None' condition) marked with asterisks $p<3e^{-5}$ (Wilcoxon test with Bonferroni correction). **b)** Differences in speed magnify the code for all three variables (Context, Direction, Position), as can be seen by the difference in decoding for the original spike data (Orig.) and the speed-equalized data (SE). The SE data is the same as shown in Figure 4b. Orig. vs SE: $p<6e^{-8}$ for all three variables, Wilcoxon test. **c)** Comparison of context decoding shows that using positions inferred by an SVM instead of real position in the maze still yields the advantages of conditional decoding. This shows that if the brain decodes these variables sequentially the benefits remain. This analysis was done on the original PVs, calculated on raw spike data (speed and direction filtered). The first two box plots are calculated the same way as in Figure 4d. Real position means that the conditional variable was the real position values from the behavior data. Inferred position refers to the position labels estimated by a Global positional decoder (see Extended Methods). Global decoding from joint labels means that position and context were combined into a single label for training and testing the SVM. The accuracy we report is only whether predicted and actual labels match in terms of context. All pairs $p<0.01$ two-sided paired t-test with Bonferroni correction. **d)** Same as Figure 5c, but for position decoding: cross-validated decoding accuracy shown as a function of the fraction of PCs used for decoding, normalized by the accuracy each decoder achieved on the complete dataset (using all PCs). The relative accuracies of global (dashed) and conditional (solid, conditional on context) decoders were highly overlapping in all learning stages, which indicates that training did not affect the fraction of neural activity variance devoted to position coding, neither globally nor in a context-specific manner. Shaded areas around the lines = SEM over all sessions in each behavior group, n=8, 9 and 8 for each group, respectively. Vertical dotted lines show the fraction of PCs needed by the global and conditional Position decoders to reach 90% of their respective maximum accuracy.

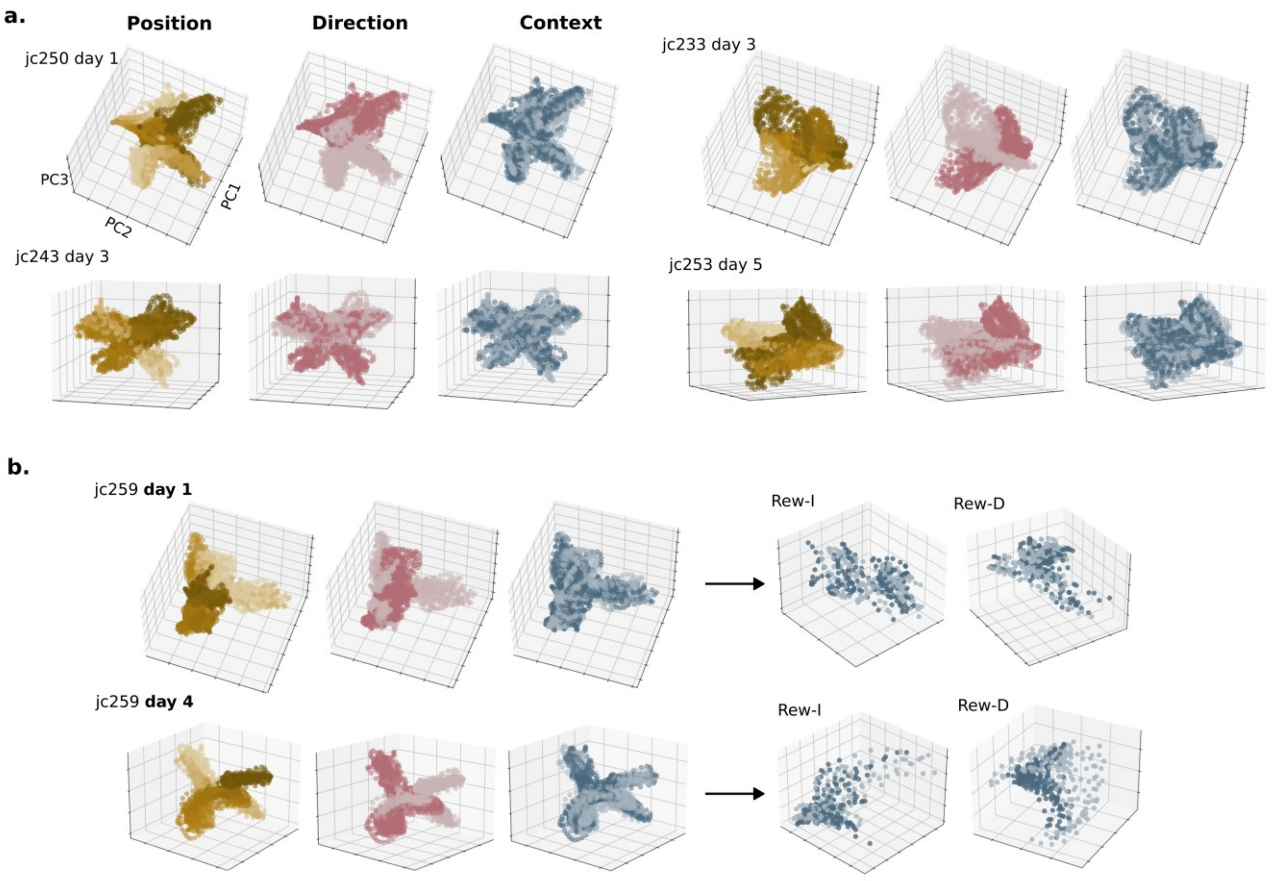


**Figure S6. PCA projections of neural data.** Example projections of the population vectors from different animals/sessions onto the first 3 principal components. **a)** At all stages of learning, Positions were well separated in the first principal components, as well as the two running directions. The separation was less clear for the Context variable. Shades of yellow indicate different maze Positions, the shades of blue indicate the two Contexts and shades of pink indicate the two movement Directions**. b)** Two extra examples comparing day 1 (Naive) and day 4 (Expert) for one specific animal (jc259), with a zoom into the data points for specific reward locations, showing a separation of contexts in the first PCs happened only at Rew-D in the Expert session, but not at Rew-I.
